# A Mechanosensitive GPCR at vestibular kinocilium is required for normal balance

**DOI:** 10.64898/2025.12.06.692705

**Authors:** Shu-Hua Zhou, Xiao-Hui Wang, Qi-Yue Zhang, Jia-Ning Xu, Jia-Le Wang, Wei-Feng Zhang, Jin-Hui Ding, Ming-Wei Wang, Ya-Qi Wang, Peng-Han, Ren-Jie Chai, Wen-Wen Liu, Xiao Yu, Yu Sun, Zhao Yang, Jin-Peng Sun

**Author notes:** Correspondence: Jin-Peng Sun, Zhao Yang, Yu Sun, Xiao Yu.

## Abstract

Intracellular calcium increase and neurotransmitter release in vestibular hair cells (VHCs) play central roles in equilibrioception, which is one of the basic senses essential for daily life activities and movement in mammals. Independent of mechano-electrical transduction (MET), whether Gq protein-coupled receptor (GqPCR) signaling participate in the regulation of intracellular calcium dynamics and induce neuronal transmitter release in hair cells remains unknown. We screened mechanosensitive GqPCRs in VHCs and found that a Class C GPCR, metabotropic glutamate receptor 2 (mGlu2), is expressed in kinocilia and is essential for normal balance. Notably, the dispensable role of mGlu2 in normal hearing is consistent with absent of mature kinocilia in cochlea hair cells. Different from the conventional mGlu2-Gi signaling, the sensing of mechanical signals by mGlu2 activates the Gq‒PLCD4 pathway, increases intracellular calcium concentration and promotes neurotransmitter release in VHCs. Hair cell-specific deficiency of either *Grm2* or *Gnaq*, or knockdown of *Plcd4* expression, but not deficiency of another GqPCR *Gpr68*, causes significant balance deficits. Reintroduction of mGlu2 into the VHCs of *Pou4f3-CreER^+/−^ Grm2^fl/fl^* mice restore vestibular functions. Our study reveals a previously uncharacterized role of GPCR signaling in equilibrioception and provides important insight into kinocilia signaling in VHCs, which are absent in mature cochlear hair cells.

## Introduction

Sensory information about motion, balance, and spatial orientation is provided primarily by the peripheral vestibular system in the inner ear, which includes two otolith organs (the utricle and saccule) and three semicircular canals (the anterior, posterior and horizontal canals) that regulate the sensation of linear acceleration and rotational head movements, respectively. Dysfunction of the vestibular system due to genetic or environmental causes leads to balance disorders, which affect approximately 30% of the general population, resulting in reduced quality of life and imposing a significant socioeconomic burden^1,2^. Equilibrioception is achieved primarily by the activation of vestibular hair cells (VHCs) through mechanoelectrical transduction (MET), which transforms head motion or tilt-evoked mechanical stimuli into electrical signals^3^. Changes in the membrane potential promote the entry of Ca^2+^ into cells through the opening of presynaptic voltage-gated Ca^2+^ channels at the basolateral membrane of VHCs, which triggers neurotransmitter (mainly glutamate) release to afferent neurons and conveys positional information to the central nervous system^4^. In addition to voltage-gated Ca^2+^ influx, Ca^2+^ levels in VHCs are also regulated by calcium-induced calcium release (CICR) from presynaptic Ca^2+^ stores through the opening of Ca^2+^-permeable channels such as the ryanodine receptor (RyR), which together contribute to transmitter exocytosis^5,6^.

In addition to Ca^2+^ channels, intracellular Ca^2+^ homeostasis can also be regulated by G protein-coupled receptors (GPCRs), especially Gq protein-coupled receptors (GqPCRs), which activate phospholipase C (PLC), an enzyme that hydrolyzes phosphatidylinositol phosphate to diacylglycerol and inositol trisphosphate, leading to intracellular Ca^2+^ mobilization^6^. In addition to responding to chemical cues, several GPCRs can also sense force and translate mechanical stimuli into Ca^2+^ signals. For example, whereas the histamine H1 receptor expressed in endothelial cells induces Ca^2+^ transients by sensing fluid shear stress and regulating flow-induced vasodilation, the adhesion G-protein coupled receptor ADGRE2 is thought to act as a mechanosensor in mast cells to promote mast cell degranulation through enhanced Ca^2+^ signaling^7–9^. Importantly, whether and how mechanosensitive GPCRs contribute to the Ca^2+^ homeostasis in VHCs remain largely unexplored.

In the present study, we screened the mechanosensitive potential of GPCRs in utricular hair cells in the activation of Gq signaling. We identified a receptor, mGlu2, which is expressed on kinocilia, as a mechanosensitive GqPCR that is required for normal balance. Global knockout (KO) or inducible hair cell-specific KO of *Grm2* (the gene encoding mGlu2) impaired balance in mice. Genetic ablation of *Grm2* did not affect MET response in VHCs but impaired the glutamate release and intracellular Ca^2+^ response. Further mechanistic studies revealed that mGlu2 converts mechanical stimuli into Ca^2+^ mobilization through the activation of the Gq‒PLCD4‒IP3R pathway. Collectively, our findings suggest that a mechanosensitive GPCR expressed on the kinocilia of VHCs is required for normal balance through a previously uncharacterized mechanochemotransduction mechanism.

## Results

### Deficiency of the mechanosensitive GqPCR mGlu2 impairs balance but not hearing function

Single-cell RNA sequencing (scRNA-seq) data indicate that many GPCRs are expressed in VHCs^10^. We speculate that a mechanosensitive GqPCR in VHCs, which translates mechanical stimuli into an intracellular Ca^2+^ response, may contribute to vestibular functions (Figure 1A). We therefore examined the mechanosensitive potential of the 35 most highly expressed GPCRs in mouse utricular hair cells by leveraging a magnetic-bead-based mechanical stimulation assay ^11,12^. We found that mGlu2 and GPR68 are the only two receptors that induce Gq signaling in response to mechanical stimulation at 10 pN (Figures 1B and 1C). Further signaling pathway profiling using BRET-based G protein dissociation assays revealed that mechanical forces only activate the Gq pathway, but not other pathways such as Gs, Gi, and G12, downstream of mGlu2 or GPR68 (Figure 1D). The EC50 values of force-induced mGlu2 and GPR68 activation were measured as 6.5 ± 0.2 pN and 4.6 ± 0.1 pN, respectively (Figure 1D). As a negative control, the untargeted force induced by polylysine-coated paramagnetic beads (Ctrl beads) did not induce the activation of any tested G protein subtypes in GPR68- or mGlu2-expressing HEK293 cells, suggesting the specific mechanosensitivity of GPR68 and mGlu2 (Figures 1D, S1A and S1B). In contrast to the force-induced mGlu2 activation, the glutamate, a known endogenous agonist of mGlu2, only elicited Gi signaling in mGlu2-overexpressed HEK293 cells (Figures S1C and S1D).

**Fig. 1.**
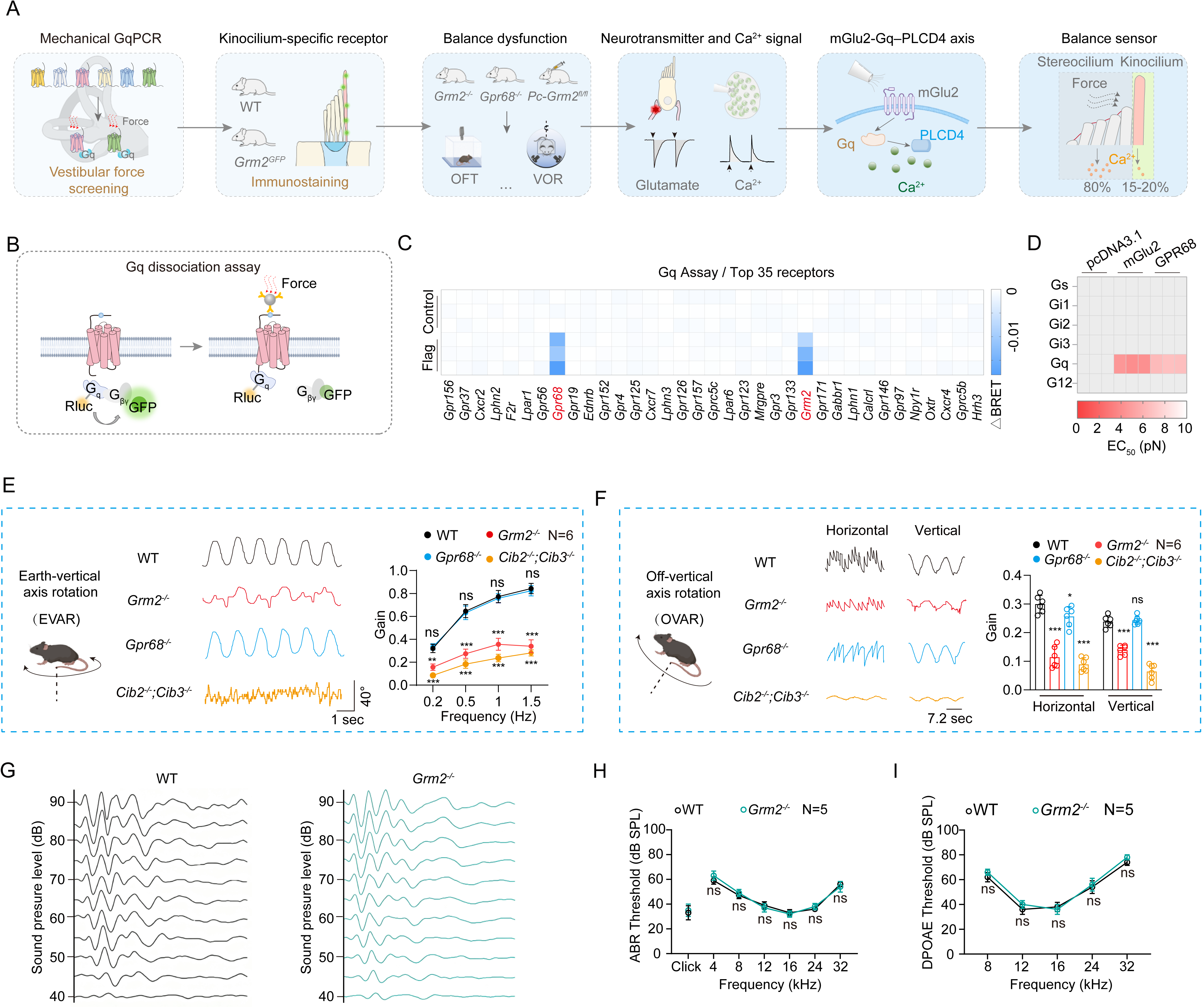
Screening of Gq-coupled mechanosensitive GPCRs in vestibular hair cells. **(A)** Schematic diagram showing the research flow for identifying mGlu2 as a mechanosensitive Gq-coupled GPCR (GqPCR) in utricular hair cells and its functional roles in balance regulation. **(B)** Schematic representation of the screening assay used for identifying mechanosensitive GqPCRs in HEK293 cells. Forces are applied on Flag antibody-coated magnetic beads via a magnetic system, which bind to the N-terminal Flag-tagged receptor and induces receptor activation. **(C)** Summary of the force-induced Gq activation downstream of the top 35 abundantly-expressed GPCRs in mouse utricular hair cells. A force of 10 pN was applied to the receptors and the Gq activation was measured by a G protein dissociation BRET assay, which was presented as a heatmap (n = 3). **(D)** Summary of the potency of force-induced G protein subtypes activation in mGlu2*-* or GPR68-overexpressed HEK293 cells (n=3). **(E)** Representative recording curves (left) and quantification (right) of the VOR gain responses of WT, *Grm2^−/−^*, *Gpr68^−/−^* and *Cib2^−/−^; Cib3^−/−^* mice to earth-vertical axis rotation (0.2-1.5 Hz, 40°/s peak velocity sinusoidal, whole-body passive rotation, N = 6 mice per group). **(F)** Representative recording curves (left) and quantification (right) of WT, *Grm2^−/−^*, *Gpr68^−/−^*and *Cib2^−/−^; Cib3^−/−^* mice to off-vertical axis rotation (50°/s, whole-body passive rotation, N = 6 mice per group). **(G-I)** ABR waveforms at click (G), ABR thresholds (H), and DPOAE thresholds (I) of WT and *Grm2^−/−^* mice at P30 (N = 5 mice per group). **(E, F, H, I)** *P < 0.05; **P < 0.01; ***P < 0.001; ns, no significant difference. Gene knockout mice compared with WT mice. The bars indicate mean ± SEM values. Data were statistically analyzed using one-way **(F)** or two-way **(E, H, I)** ANOVA with Dunnett’s post hoc test.

We then investigated the potential vestibular functions of these two GPCRs by comparing the performance of *Grm2*^−/−^ mice and *Gpr68*^−/−^ mice with that of their wild-type (WT) littermates in equilibration-related tests. *Cib2*^−/−^; *Cib3*^−/−^ mice, which have significant balance defects, were used as positive controls^13^. The deficiency of *Grm2* or *Gpr68* in the corresponding gene KO mice was verified by genotyping PCR and western blotting (Figures S1E-S1L). The KO mice were viable and had body weights comparable to those of their WT littermates (Figure S1M). The vestibular functions of the mice at P40 were assessed through the open field test. Compared with their WT littermates, *Grm2*^−/−^ mice displayed significant decreases in locomotion and circling (Figures S2A-S2C). Moreover, the *Grm2*^−/−^ mice also exhibited poorer swimming performance than their WT littermates (Figure S2D). In contrast, the *Gpr68*^−/−^ mice behaved normally in both tests (Figures S2A-S2D). We then recorded the vestibuloocular reflexes (VORs) of the P40 mice in response to sinusoidal head rotations. Compared with WT mice, *Grm2*^−/−^ mice exhibited an approximately 30%-60% decrease in compensatory VOR gain in response to earth-vertical or off-vertical axis rotations (Figures 1E and 1F). In contrast, *Gpr68*^−/−^ mice presented only subtle abnormalities in the response to off-vertical axis rotation, with an approximately 15% reduction in horizontal compensatory VOR gain (Figures 1E and 1F). These results suggest that mGlu2 plays a more important role than *Gpr68* in balance maintenance. Intriguingly, despite its important regulatory role in balance, mGlu2 seemed to be dispensable for hearing since *Grm2*^−/−^ mice presented normal auditory brainstem response (ABR) thresholds and distortion product otoacoustic emission (DPOAE) values at different testing frequencies, which were comparable to those of WT mice (Figures 1G-1I).

### Expression of mGlu2 in utricular hair cells

Our quantitative RT-PCR result showed a dynamic pattern of mGlu2 expression in utricular hair cells, with the highest mRNA levels detected at around P30 (Figures S3A). We next examined the expression pattern of mGlu2 in the mouse utricle via whole-mount immunostaining using an anti-mGlu2 antibody, the specificity of which was supported by both *in vitro* and *in vivo* western blot analysis (Figures S1F, S1G and S3B). Immunostaining for mGlu2 was detected in approximately 90% of utricular hair cells from WT mice at P40 (Figures 2A-2C). Specifically, mGlu2 immunofluorescent puncta were observed along the kinocilia and were colocalized with α-tubulin, which was absent in *Grm2^−/−^*mice of the same age (Figures 2A-2C). The specific expression of mGlu2 in the kinocilia of utricular hair cells was further supported by the usage of a *Grm2*^GFP^ transgenic knock-in mouse line, which was generated by fusing a GFP fluorophore to the C-terminus of mGlu2 (Figures 2D-2F). We further explored the subcellular distribution of mGlu2 in utricular hair cells via optical sectioning microscopy, which again supported the expression of mGlu2 in kinocilia but not in stereocilia or at the apical surface (Figure 2G). Notably, despite its absence in supporting cells, mGlu2 was also expressed at the interface between hair cells and afferent neurons, as revealed by its colocalization with Tuj1, which is consistent with a previous report and suggests a regulatory role for mGlu2 in neurotransmitter release^14^ (Figure S3C).

**Fig. 2.**
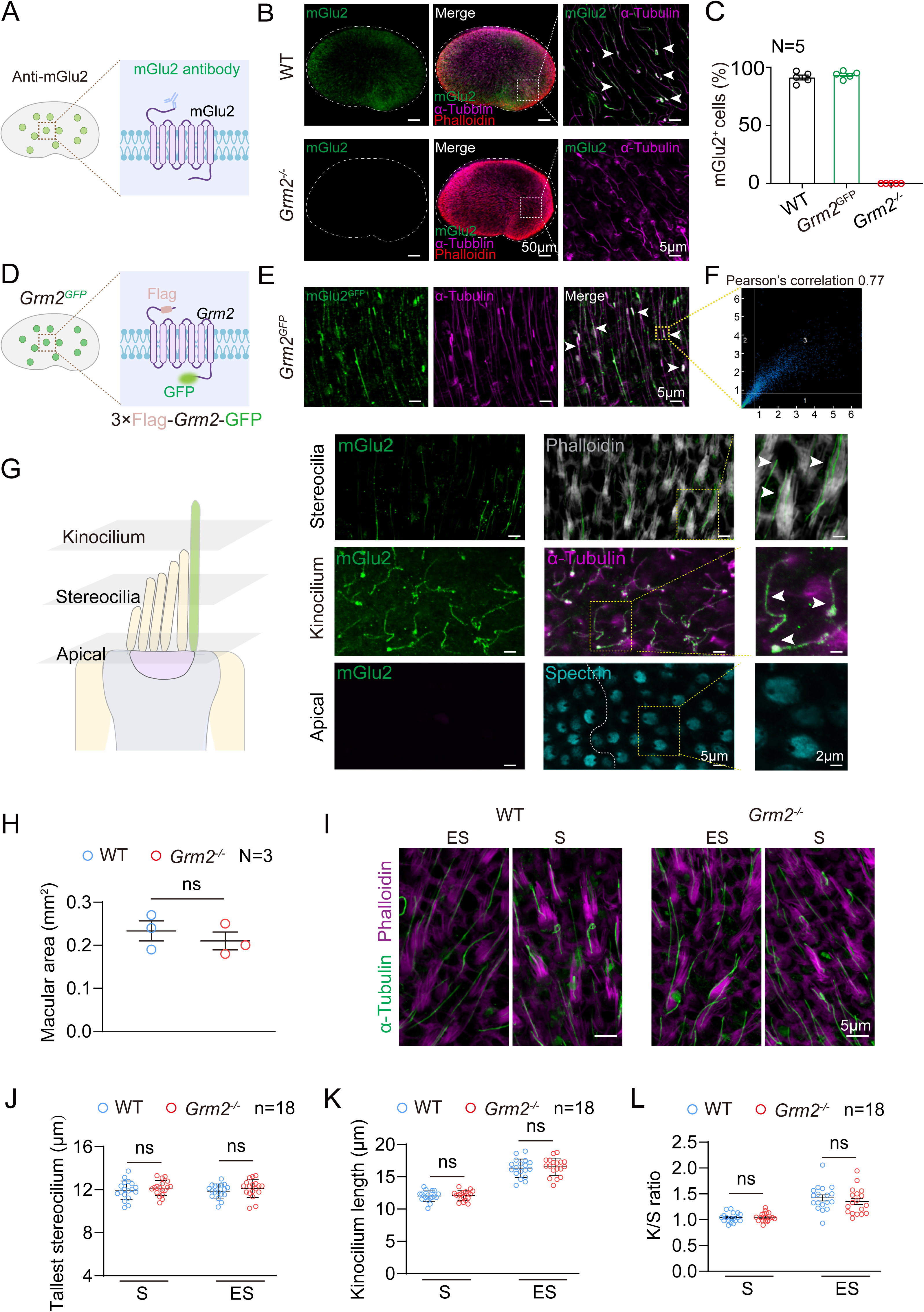
mGlu2 is expressed at the kinocilium of utricular hair cells. **(A)** Schematic diagram showing the whole-mount immunostaining of mGlu2 expression in mouse utricles using an anti-mGlu2 antibody. **(B)** Coimmunostaining of mGlu2 (green) with phalloidin (red) and α-tubulin (magenta) in utricle whole mounts derived from WT mice and *Grm2^−/−^*mice (N = 5 mice per group). Scale bar, 50 μm and 5 μm for low- and high-magnification views, respectively. **(C)** Quantitative analysis of mGlu2 expression in utricular hair cells from WT mice, *Grm2^GFP^* mice or *Grm2^−/−^* mice (N = 5 mice per group). **(D)** Schematic diagram showing the whole-mount immunostaining of mGlu2 expression in the utricles derived from *Grm2^GFP^* mice. **(E-F)** Expression of mGlu2-GFP (green) with α-tubulin (magenta) in utricle whole mounts of P40 mice. Arrows indicate coimmunostaining of mGlu2 with α-tubulin. Scale bar, 5 μm. A Pearson’s correlation analysis **(F)** of the fluorescence intensities of mGlu2 and α-tubulin at the kinocilium was performed, revealing a correlation coefficient of 0.77. **(G)** Left panel: Diagram of utricular hair cells showing the selected optical planes (kinocilium, stereocilia or apical surface) for imaging by confocal microscopy. Right panel: Coimmunostaining of mGlu2 (green) with spectrin (cyan), phalloidin (gray) or α-tubulin (magenta) at different optical planes of utricular hair cells in P40 mice. Arrows indicate coimmunostaining of mGlu2 with kinocilium. Scale bar, 5 μm and 2 μm for low- and high-magnification views, respectively. **(H)** Quantification of the size of utricles derived from WT and *Grm2^−/−^* mice (N = 3 mice per group). **(I)** Immunostaining of kinocilium (labeled with α-tubulin, green) and stereocilia (labeled with phalloidin, magenta) in utricle whole mounts derived from WT and *Grm2^−/−^* mice (N = 3 mice per group). Scale bar, 5 μm. **(J)** Quantification of the length of tallest stereocilia in S (striolar) or ES (extrastriolar) region of utricles derived from WT and *Grm2^−/−^* mice. Data are correlated to Fig. 2I (n = 18 hair cells from 3 mice per group). **(K, L)** Quantification of the length of kinocilium **(K)** and the ratio of lengths of the kinocilium to tallest stereocilia **(L)** in S or ES region of utricle whole mounts derived from WT and *Grm2^/-^* mice. Data are correlated to Fig. 2I (n = 18 hair cells from 3 mice per group). **(H, J-L)** ns, no significant difference. *Grm2^−/−^* mice compared with WT mice. The bars indicate mean ± SEM values. All data were statistically analyzed using unpaired two-sided Student’s *t* test.

To investigate whether the balance deficits of *Grm2^−/−^*mice were due to developmental defects, we next investigated the morphological and anatomical features of the utricles in *Grm2*^−/−^ mice at P40. The macula size in the *Grm2*^−/−^ mice was comparable to that in the WT mice (Figures 2H and S3D). Moreover, the kinocilium length and the K/S ratio (the ratio of the length of the kinocilium to the height of the tallest stereocilium), which are important indicators of utricular physiology^15^, were comparable between the utricles of *Grm2^−/−^* mice and those of their WT counterparts (Figures 2I-2L). Therefore, these data suggest that the morphological integrity of the utricle was maintained in *Grm2*^−/−^ mice, at least at P40.

### mGlu2 expressed in VHCs is required for balance

mGlu2 is expressed throughout the central nervous system, where it actively participates in the regulation of cognition and emotion by modulating glutamatergic activity at synapses in the brain^16,17^. To evaluate the specific functions of mGlu2 in VHCs, we generated *Pou4f3-CreER^+/−^*; *Grm2^fl/fl^* mice, which enabled tamoxifen-induced hair cell-specific ablation of mGlu2 (Figures 3A, S4A and S4B). *Pou4f3* is a hair cell-specific transcription factor downstream of ATOH1, and scRNA-seq data (GSE71982) have indicated that *Pou4f3* is expressed in all mGlu2-positive utricular hair cells^10^. A decrease in mGlu2 expression in the utricles of *Pou4f3-CreER^+/−^*; *Grm2^fl/fl^* mice was verified by western blot analysis (Figures S4C and S4D). Moreover, mGlu2 immunostaining on kinocilia was specifically ablated in the utricular hair cells of *Pou4f3-CreER^+/−^*; *Grm2^fl/fl^* mice but not in their *Pou4f3-CreER^+/−^*; *Grm2^+/+^* counterparts (Figures S4E and S4F). In contrast, the expression of mGlu2 in other regions, such as the vestibular nuclei in the brainstem, was not significantly affected in *Pou4f3-CreER^+/−^*; *Grm2^fl/fl^* mice (Figures S4G and S4H).

**Fig. 3.**
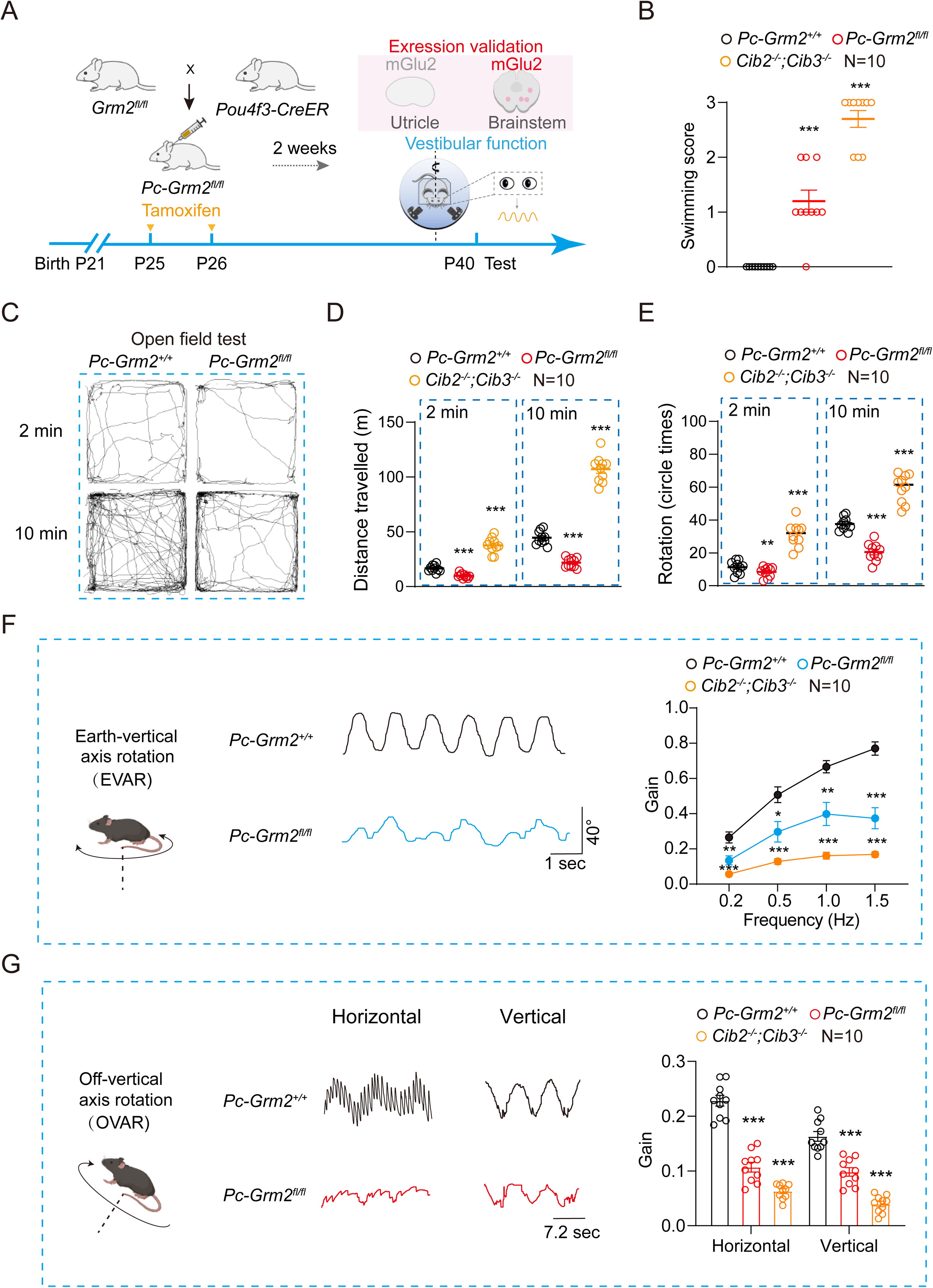
Conditional knockout of *Grm2* in mouse hair cells impairs balance. **(A)** Schematic representation of the crossbreeding strategy to generate hair cell-specific *Grm2* knockout mice and the time scales for vestibular functional analysis. The *Pou4f3-CreER^+/−^*; *Grm2^fl/fl^* mice (referred to as *Pc-Grm2^fl/fl^*) or *Pou4f3-CreER^+/−^*; *Grm2^+/+^* mice (referred to as *Pc-Grm2^+/+^*) were treated with tamoxifen (75 mg/kg) dissolved in corn oil through round window membrane injection at P25 (left ear) and P26 (right ear) consecutively, and the vestibular behavior tests were performed at P40. **(B-E)** Quantification of the swimming scores (**B**), representative tracks (**C**), travelling (**D**) and circling activity (**E**) in the open field test of *Pc-Grm2^fl/fl^*mice, *Pc-Grm2^+/+^* mice and *Cib2^−/−^; Cib3^−/−^* mice (N = 10 mice per group). **(F)** Representative recording curves (left) and quantification of the VOR gain values (right) of *Pc-Grm2^fl/fl^* mice, *Pc-Grm2^+/+^*mice and *Cib2^−/−^; Cib3^−/−^* mice in response to earth-vertical axis rotation (N = 10 mice per group). **(G)** Representative recording curves (left) and quantification of the VOR gain values (right) of *Pc-Grm2^fl/fl^*mice, *Pc-Grm2^+/+^* mice and *Cib2^−/−^; Cib3^−/−^* mice in response to off-vertical axis rotation (N = 10 mice per group). (**B, D-G**) *P < 0.05; **P < 0.01; ***P < 0.001; ns, no significant difference. *Pc-Grm2^fl/fl^* or *Cib2^−/−^; Cib3^−/−^* mice compared with *Pc-Grm2^+/+^* mice. The bars indicate mean ± SEM values. Data were statistically analyzed using one-way (**B, D, E, G**) or two-way ANOVA (**F**) with Dunnett’s post hoc test.

We next performed a series of balance behavioral tests using *Pou4f3-CreER^+/−^*; *Grm2^fl/fl^* mice. The performance of *Pou4f3-CreER^+/−^*; *Grm2^fl/fl^* mice in the open field test was almost identical to that of *Grm2^−/−^* mice. Compared with those of the *Pou4f3-CreER^+/−^*; *Grm2^+/+^* mice, *Pou4f3-CreER^+/−^*; *Grm2^fl/fl^*mice exhibited decreased circling and locomotion in the open field test and poorer swimming performance (Figures 3B-3E). Moreover, *Pou4f3-CreER^+/−^*; *Grm2^fl/fl^* mice presented an approximately 35–50% decrease in VORs in response to both earth-vertical and off-vertical axis rotation. Similar to *Grm2^−/−^* mice, the *Pou4f3-CreER^+/−^*; *Grm2^fl/fl^* mice displayed a maximal reduction in compensatory VOR gain in response to earth-vertical rotation at a frequency of 1.5 Hz compared with their respective control littermates (Figures 3F and 3G). Despite significant balance deficits, the *Pou4f3-CreER^+/−^*; *Grm2^fl/fl^* mice presented normal utricle morphology at P40, as revealed by a macula size, hair cell density and kinocilia/stereocilia length comparable to those of *Pou4f3-CreER^+/−^*; *Grm2^+/+^*mice (Figures S5A-S5D). In addition, the core components of the MET channel complex, such as TMC1 and TMC2, exhibited normal expression levels and localization in the utricular hair cells of *Pou4f3-CreER^+/−^*; *Grm2^fl/fl^* mice (Figures S5E-S5G). Collectively, these results suggest that mGlu2 in VHCs plays an indispensable role in the regulation of balance.

### Reintroduction of *Grm2* in vestibular hair cells rescues balance of *Grm2*-deficient mice

To further assess the specific contribution of VHC-expressed mGlu2, we reintroduced mGlu2 expression into the utricular hair cells of *Pou4f3-CreER^+/−^*; *Grm2^fl/fl^*mice via AAV strategy (Figure 4A). An AAV-ie vector was selected, and the promoter region was substituted with that of the *Grm2* gene (1000 bp). The *Grm2* cDNA sequence encoding a mGlu2 fragment with a truncation of the N-terminal 555 residues was packaged in the AAV-ie vector due to the limited capacity of the vector (<3.5 kb) (Figure S6A). This truncated version of mGlu2 (mGlu2^Δ1–555^) showed comparable Gq activity as the intact WT mGlu2 in response to mechanical stimulation but was insensitive to glutamate (Figures 4B, S6B and S6C). The detailed structural and molecular mechanism of force-induced mGlu2 activation has been described in details in our parallel manuscript. To specifically label mGlu2 expression in VHCs, an EGFP was fused to the C-terminus of mGlu2^Δ1–555^. This virus, referred to as AAV-ie-mGlu2, was delivered into P3 *Pou4f3-CreER^+/−^*; *Grm2^fl/fl^* mice through round window membrane injection, and the AAV-ie-*Grm2pr*-EGFP vector, which only has *Grm2* promoter but no encoding sequence of mGlu2, was used as a negative control. 14 days after the virus administration, mGlu2 expression was detected in the kinocilia of utricular hair cells, but not in the brainstem, of the AAV-ie-mGlu2-treated *Pou4f3-CreER^+/−^*; *Grm2^fl/fl^* mice, as shown by western blotting analysis (Figure S6D and S6E). In *Pou4f3-CreER^+/−^*; *Grm2^fl/fl^* mice injected with AAV-ie-*Grm2pr*-EGFP, the utricular hair cells were labeled with EGFP, but no mGlu2 expression in the kinocilia was detectable (Figure S6F).

**Fig. 4.**
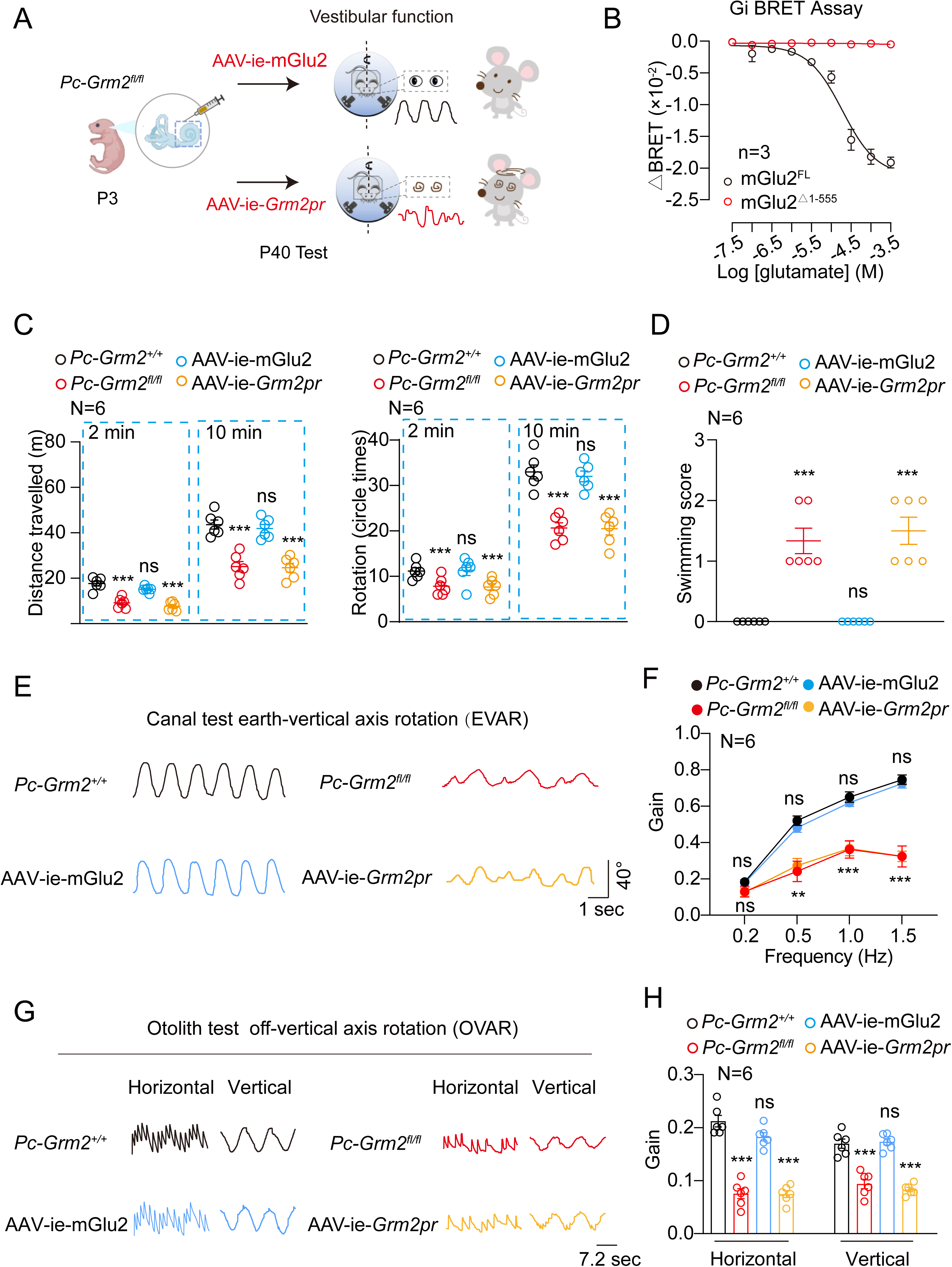
Re-expression of mGlu2 in hair cells of *Grm2*-deficient mice rescues the vestibular function. **(A)** Schematic representation of the strategy to reintroduced mGlu2 expression in the hair cells of the *Pc-Grm2^fl/fl^* mice via an AAV-ie strategy and the vestibular function validation. **(B)** Dose-dependent Gi activation in HEK293 cells transfected with full length mGlu2 (mGlu2^FL^) or mGlu2^Δ1–555^ in response to glutamate stimulation. Representative curves from three independent experiments are shown (n=3). **(C)** Quantification of the travelling (left) and circling (right) activity of *Pc-Grm2^+/+^*, *Pc-Grm2^fl/fl^*, AAV-ie-mGlu2 mice and AAV-ie-*Grm2pr* mice in the open-field test (N = 6 mice per group). **(D)** Quantification of the swimming scores of *Pc-Grm2^+/+^*, *Pc-Grm2^fl/fl^*, AAV-ie-mGlu2 mice and AAV-ie-*Grm2pr* mice (N = 6 mice per group). **(E-H)** Representative recording curves (left) and quantification of the VOR gain responses (right) of *Pc-Grm2^+/+^*, *Pc-Grm2^fl/fl^*, AAV-ie-mGlu2 mice and AAV-ie-*Grm2pr* mice to earth-vertical axis (**E, F**) or off-vertical axis (**G, H)** rotation (N = 6 mice per group). (**C, D, F, H**) **P < 0.01; ***P < 0.001; ns, no significant difference. *Pc-Grm2^fl/fl^*, AAV-ie-mGlu2 mice or AAV-ie-*Grm2pr* mice compared with *Pc-Grm2^+/+^* mice. The bars indicate mean ± SEM values. Data were statistically analyzed using one-way (**C, D, F**) or two-way (**F**) ANOVA with Dunnett’s post hoc test.

We next examined the balance-related behaviors of the mice. Notably, *Grm2*-deficient mice treated with AAV-ie-mGlu2 exhibited significantly improved performance in the open field test and forced swimming test, with their performance being comparable to that of the *Pou4f3-CreER^+/−^*; *Grm2^+/+^*mice (Figures 4C and 4D). Moreover, after AAV-ie-mGlu2 injection, the VORs of *Pou4f3-CreER^+/−^; Grm2^fl/fl^* mice in response to both earth-vertical and off-vertical axis rotation recovered to levels comparable to those of WT mice (Figures 4E-4H). In contrast, *Grm2*-deficient mice administered with the control AAV-ie-*Grm2pr*-EGFP showed no significant improvement in any of the behavior tests or in VORs. These findings further support that mGlu2 expressed in VHCs specifically contributes to normal balance.

### Mechanosensation by mGlu2 induces neurotransmitter release and Ca^2+^ response in utricular hair cells in a MET independent manner

Signals related to the bending of stereociliary bundles in response to mechanical stimulation are transmitted from VHCs to vestibular neurons via the release of neurotransmitters (Figure 5A). We examined whether the loss of mGlu2 in the VHCs affects force-induced neurotransmitter release via expressing the glutamate reporter R^ncp^-iGluSnFR in vestibular afferent neurons^18^ (Figure 5B). A fluid jet was applied to the hair bundles of isolated utricular explants to mimic physiological stimulation *in vivo*. Fluid jet-induced glutamate release from WT utricular hair cells was readily detected, as revealed by a decrease in the fluorescent density of the glutamate reporter (Figure 5C). Compared with their WT counterparts, utricular hair cells derived from *Pou4f3-CreER^+/−^; Grm2^fl/fl^* mice presented an approximately 45% decrease in glutamate release in response to fluid jet stimulation (Figures 5C and 5D). In addition, the recovery time constant of glutamate release in *Grm2*-deficient utricular hair cells, when fitted to a monoexponential equation, was significantly lower than that in WT utricular hair cells (Figure S7A). Using the same stimulation system, we further assessed the Ca^2+^ response in utricular hair cells preloaded with the Fura-2 AM probe (Figure 5B). Consistent with the glutamate release results, the fluid jet-stimulated Ca^2+^ response (area under the curve) of hair cells from *Pou4f3-CreER^+/−^Grm2^fl/fl^* mice was approximately 40% lower than that of hair cells from their *Pou4f3-CreER^+/−^Grm2^+/+^* littermates (Figures 5E and 5F). The peak value of the Ca^2+^ response in single utricular hair cells from *Pou4f3-CreER^+/−^Grm2^f.l/fl^* mice was approximately 85% of that in control cells (Figure S7B). These data suggest that mGlu2 plays an important role in neurotransmitter release and Ca^2+^ signaling in utricular hair cells in response to mechanical stimulation.

**Fig. 5.**
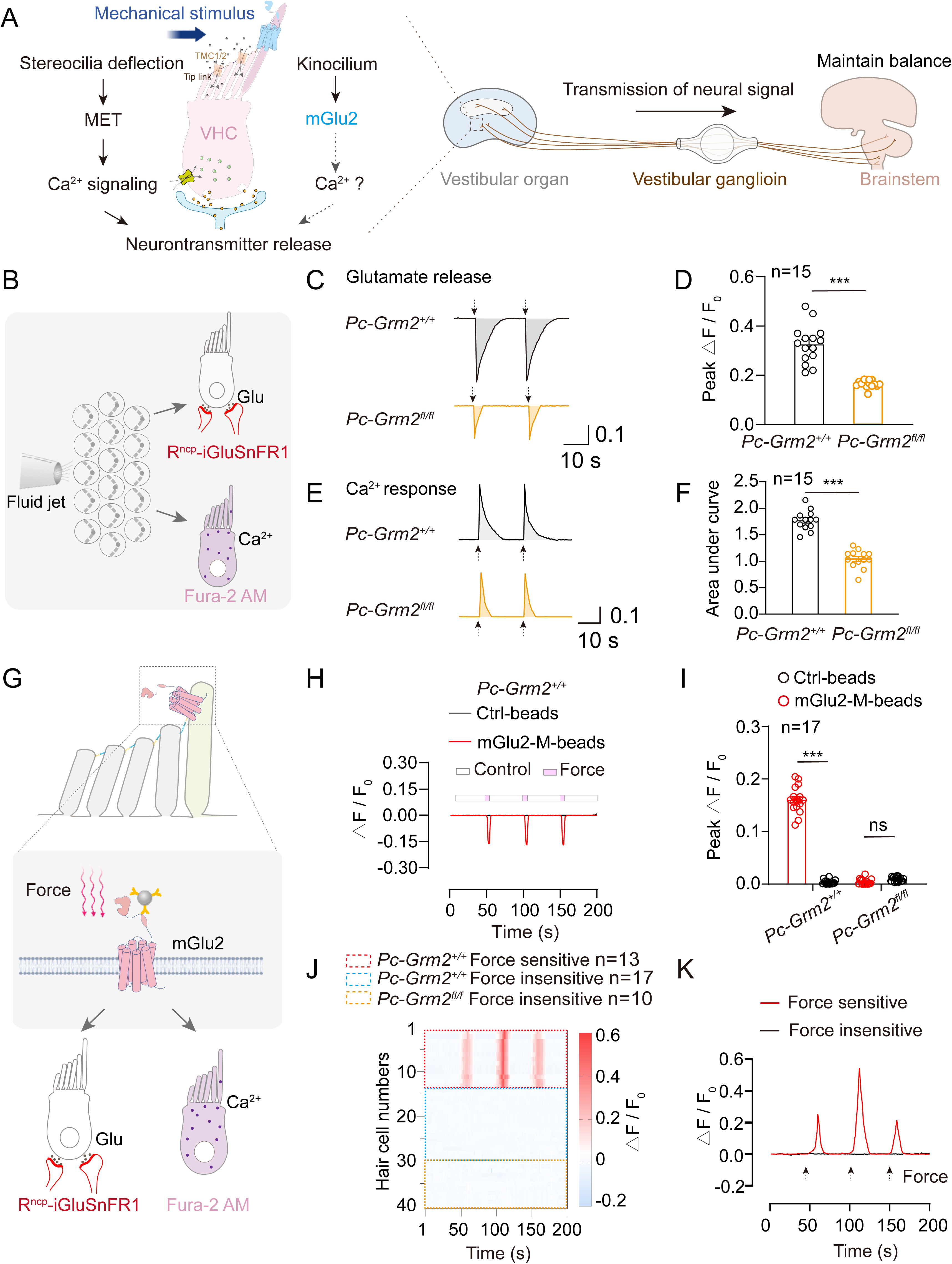
Force sensation by mGlu2 induces glutamate release and Ca^2+^ signals in utricular hair cells. **(A)** Schematic diagram showing the process of Ca^2+^ signaling, neurotransmitter release and neural electrical signal from vestibule organ to central system after the activation of hair cells by mechanical stimulus. **(B)** Schematic diagram showing the detection of fluid jet-stimulated glutamate release by a glutamate reporter R^ncp^-iGluSnFR and the detection of Ca^2+^ response by Fura-2 AM in utricular hair cells. **(C, D)** Representative traces (**C**) and quantitative analysis (**D**) of fluid jet-stimulated glutamate secretion from utricular hair cells derived from *Pc-Grm2^+/+^* mice (black) or *Pc-Grm2^fl/fl^* mice (orange) at P20 (n=15). The magnitude of the glutamate secretion was characterized by ΔF/F_0_. **(E, F)** Representative traces (**E**) and quantitative analysis (**F**) of fluid jet-stimulated Ca^2+^ signals in utricular hair cells derived from *Pc-Grm2^+/+^* mice (black) or *Pc-Grm2^fl/fl^* mice (orange) at P20 (n=15). The magnitude of the Ca^2+^ response was characterized by ΔF/F_0_. **(G)** Schematic diagram showing the glutamate release and Ca^2+^ response in mouse utricular hair cells in response to force stimulation applied through magnetic beads. **(H, I)** Representative traces **(H)** and quantitative analysis **(I)** of glutamate secretion from individual utricular hair cell derived from P20 *Pc-Grm2^+/+^* mice or *Pc-Grm2^fl/fl^* mice in response to force applied through mGlu2-M-beads or Ctrl-beads (n = 17). **(J)** Heatmaps showing the Ca^2+^ responses in individual utricular hair cell derived from *Pc-Grm2^+/+^*or *Pc-Grm2^fl/fl^* mice. n=13, 17 and 10 for *Pc-Grm2^+/+^* force-sensitive cells (red box), *Pc-Grm2^+/+^* force-insensitive cells (blue box) and *Pc-Grm2^fl/fl^*force-insensitive cells (orange box), respectively. The color intensity represents the magnitude of the calcium response characterized by ΔF/F_0_. **(K)** Representative Ca^2+^ traces recorded in individual utricular hair cells derived from force-sensitive cells in *Pc-Grm2^+/+^*mice (red) or force-insensitive cells in *Pc-Grm2^fl/fl^* mice (black). **(D, F)** ***P < 0.001; ns, no significant difference. *Pc-Grm2^+/+^* mice compared with *Pc-Grm2^fl/fl^*mice. **(I)** ***P < 0.001; ns, no significant difference. Utricle explants treated with mGlu2-M-beads compared with those treated with Ctrl-beads. The bars indicate mean ± SEM values. All data were statistically analyzed using unpaired two-sided Student’s t test.

To directly evaluate the contribution of the mechanosensitive mGlu2 to glutamate release and Ca^2+^ signals, we employed a magnetic bead-based mechanical stimulation system and applied mechanical force to the utricular epithelium with mGlu2-M-beads (Figure 5G). Notably, approximately 42% of mGlu2-expressing utricular hair cells labeled with AAV-ie-*Grm2pr*-EGFP presented marked glutamate secretion in response to repeated stimulation with 10 pN force, which was abrogated in utricular hair cells from *Pou4f3-CreER^+/−^Grm2^fl/fl^*mice (Figures 5H-5I, S7C-S7E). The magnitude of mGlu2-M-bead-induced glutamate release was approximately 57% of that induced by CDH23-M-beads, which could theoretically bind to the tip link and deflect the stereocilia of VHCs to trigger the cell activation (Figures S7F and S7G). In contrast, the force applied by Ctrl-beads did not induce any detectable glutamate response (Figures S7F and S7G). Similar results were also obtained in Ca^2+^ recording of utricular hair cells, showing that repeated mechanical stimulation of the utricular epithelium via mGlu2-M-beads induced Ca^2+^ responses in approximately 44% of mGlu2-expressing hair cells but not in mGlu2-deficient hair cells (Figures 5J and 5K). The fact that the percentage of mGlu2-expressing utricular hair cells that responded to mechanical stimulation (∼44%) was lower than the percentage of mGlu2-expressing cells (∼90%) was likely due to the insufficient binding efficiency of the magnetic beads to the target protein. In support of this hypothesis, only approximately 53% of randomly selected utricular hair cells presented detectable Ca^2+^ signals in response to mechanical stimulation via CDH23-M beads (Figures S7H and S7I).

The specific expression of mGlu2 on kinocilia and the normal distribution of MET core components at stereocilia in mGlu2-deficient utricular hair cells implied that mGlu2 may be dispensable for MET. Consistent with this hypothesis, the peak MET currents of utricular hair cells from *Pou4f3-CreER^+/−^*; *Grm2^fl/fl^* mice in response to fluid jet stimulation were comparable to those of utricular hair cells from *Pou4f3-CreER^+/−^*; *Grm2^+/+^* mice (Figures 5J-5L). In contrast, compared with those of WT utricular hair cells, the peak MET currents of *Tmc1^−/−^; Tmc2^+/−^*utricular hair cells were reduced by approximately 65% (Figures S7M and S7N). Taken together, these results indicate that the force sensing by mGlu2 in VHCs induces Ca^2+^ response and neurotransmitter release in a MET-independent manner.

### Mechanosensation by mGlu2 regulates intracellular Ca^2+^ concentrations through Gq– PLCD4 pathway

mGlu2 may translate mechanical stimulation into a Ca^2+^ response in utricular hair cells by regulating Ca^2+^ influx via cation channels or by promoting Ca^2+^ release from intracellular Ca^2+^ stores. To explore the mechanism underlying the mechanical stimulation-induced increase in intracellular Ca^2+^ concentrations via mGlu2, we examined the effects of an extracellular Ca^2+^-free bath or the IP3R inhibitor 2-APB on mGlu2-M-bead-induced Ca^2+^ responses in utricular hair cells (Figures 6A). Notably, whereas the Ca^2+^ response was not significantly affected by substitution of the extracellular bath with Ca^2+^-free buffer, it was completely inhibited by pretreatment with 2-APB, thus suggesting mobilization of intracellular Ca^2+^ (Figures 6A and 6B). Moreover, the mGlu2-M-bead-induced Ca^2+^ signal was also abolished by pretreatment with Gq inhibitor YM-254890 or PLC inhibitor U73122 but not by Gs inhibitor NF449 or MET inhibitor amiloride (Figures 6A, 6B and S8A). Consistent with the observation of Ca^2+^ signals, mechanical force-stimulated glutamate release from utricular hair cells via mGlu2 activation was abrogated by pretreatment with YM-254890, U73122 or 2-APB but was unaffected by pretreatment with NF449 or MET inhibitor amiloride (Figures 6C and 6D). These data indicate that Gq signaling mediates the Ca^2+^ response downstream of mechanosensitive mGlu2 in utricular hair cells.

**Fig. 6.**
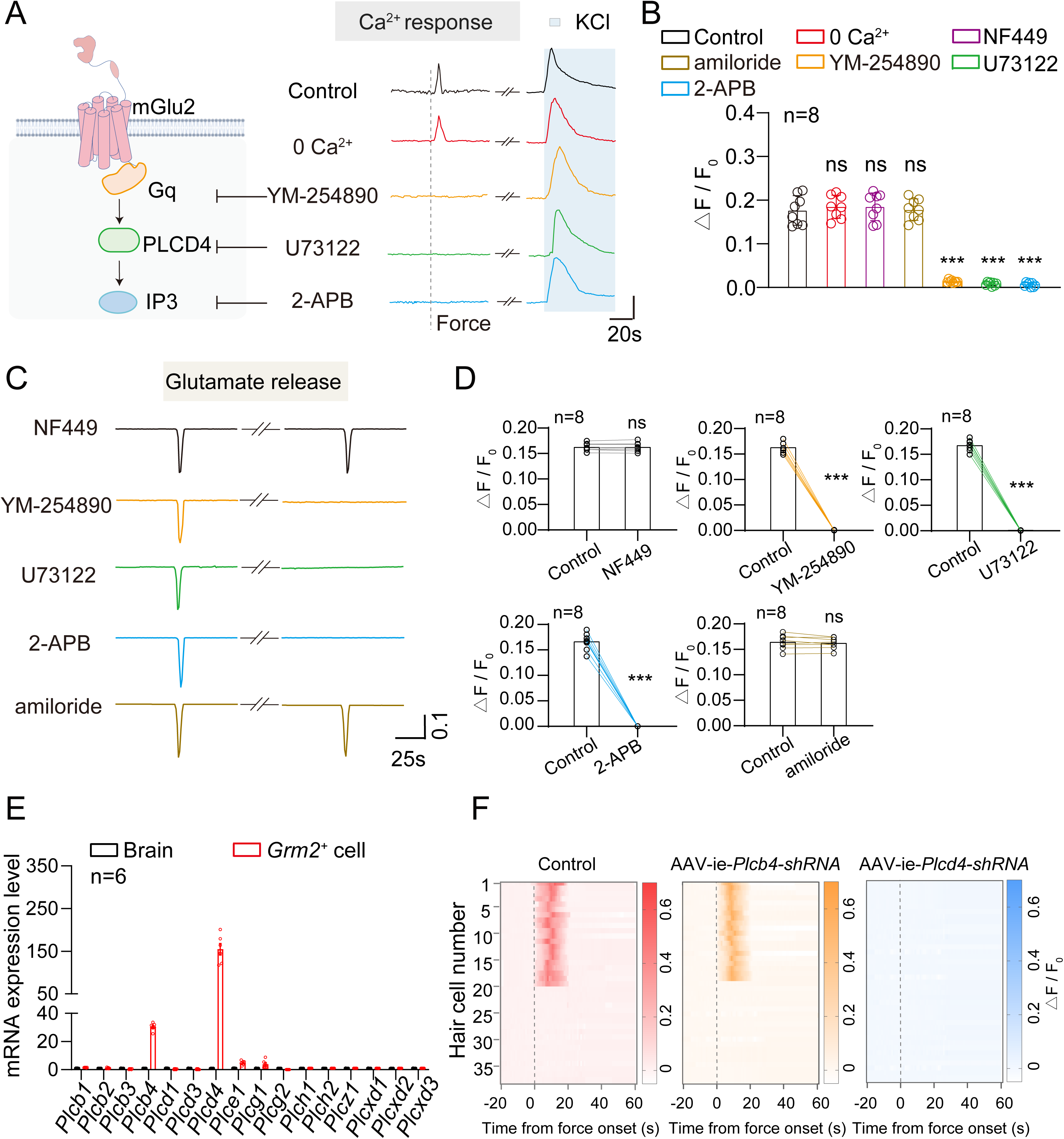
mGlu2 regulates force-stimulated Ca²⁺ response via Gq-PLCD4 pathway. **(A)** The experimental strategy to probe the Gq-PLC signaling pathway using specific inhibitor and representative traces of the effect of NF449 (20 μM), amiloride (0.2 mM), YM-254890 (5 μM), U73122(10 μM) or 2-APB (200 μM) treatment or Ca^2+^ ablation on the force-induced Ca^2+^ response in utricular hair cells labelled with AAV-ie-*Grm2pr*-EGFP (n=8 cells per group). The hair cells were stimulated with 70 mM potassium chloride to verify the Ca^2+^ response. **(B)** Quantitative analysis of the effect of NF449, amiloride, YM-254890, U73122 or 2-APB treatment or Ca^2+^ ablation on the force-induced Ca^2+^ response in utricular hair cells. **(C, D**) Representative traces **(C)** and quantitative analysis **(D)** of the effect of NF449, YM-254890, U73122, 2-APB or amiloride treatment on the force-induced glutamate secretion in utricular hair cells labelled with AAV-ie-*Grm2pr*-EGFP (n=8 cells per group). **(E)** mRNA levels of different *Plc* subtypes in mGlu2-expressing utricular hair cells derived from P10 mice measured by single-cell qRT-PCR (n = 6 hair cells). The mRNA levels of respective *Plc* subtypes in the brain were used as the reference (normalized to 1). **(F)** Heatmaps showing the Ca^2+^ responses in individual utricular hair cell derived from control, AAV-ie-*Plcb4*-*shRNA* or AAV-ie-*Plcd4*-*shRNA* treated mice in response to force stimulation applied through mGlu2-M-beads. The color intensity represents the magnitude of the calcium response, which is characterized by ΔF/F_0_. **(B, D)** ***P < 0.001; ns, no significant difference. Utricular hair cells incubated with Ca^2+^-free buffer or pretreated with NF449, amiloride, YM-254890, U73122 or 2-APB with those treated with control vehicle. The bars indicate mean ± SEM values. Data data were statistically analyzed using one-way **(B)** or paired two-sided Student’s t test **(D)**.

Since the PLC family consists of 16 members in 7 distinct subclasses, we next investigated which PLC subtype(s) participate in mGlu2-mediated Gq signaling. We assessed the expression of PLC family members within mGlu2-positive utricular hair cells using single-cell qRT‒PCR and found that *Plcb4* and *Plcd4* were the most highly expressed PLC subtypes and that their expression levels in utricular hair cells were more than 20-fold greater than those in the brain (Figure 6E). Importantly, specific knockdown of *Plcd4* in utricular hair cells nearly abolished the Ca^2+^ response induced by mechanical stimulation with mGlu2-M-beads (Figures 6F, S8B-S8E). In contrast, there was no difference in the mGlu2 bead-induced intracellular Ca^2+^ response in the *Plcb4* knockdown group compared with the control group (Figures 6F, S8B-S8E). The Ca^2+^ response induced by mechanical stimulation with CDH23-M-beads, which served as a control, was not affected by knockdown of either *Plcb4* or *Plcd4* (Figures S8D and S8F). Therefore, these results suggest that PLCD4 is an essential signaling molecule in mGlu2-mediated Gq signaling that governs Ca^2+^ mobilization in utricular hair cells.

To further verify the mechanism underlying mechanical force-stimulated mGlu2 activation, we next reconstituted the Gq-PLCD4 signaling pathway in HEK293 cells. To mimic the endogenous landscape in utricular hair cells, we adjusted the expression levels of relevant signaling components, including *Grm2*, *Gnaq* and *Plcd4*, in HEK293 cells to achieve equal expression as in native utricular hair cells (Figures S8G-S8I). We showed that both fluid jet stimulation and application of a 10 pN force via mGlu2-M-beads could induce a significant Ca^2+^ response in the reconstituted system, which was not detectable when mGlu2 was absent in the system (Figures S8J and S8K). Collectively, these results suggest that mechanical forces sensed by mGlu2 in VHCs are translated into a Ca^2+^ response and neurotransmitter release, primarily through the Gq-PLCD4 pathway.

### Gq and PLCD4 are essential for balance

To gain a deeper understanding of the regulatory role of the Gq‒PLCD4 pathway, we next detected the expression pattern of Gq and PLCD4 in utricular hair cells (Figure 7A). Whole-mount immunostaining revealed that in approximately 90% of mGlu2-positive hair cells, the Gq was expressed at the kinocilia, where it colocalized with mGlu2 (Figures 7B, 7C and S9E). Due to the lack of effective and specific antibodies targeting PLCD4, we introduced an HA-PLCD4 expression by infecting the mouse inner ear with an AAV-ie encompassing the promoter region (1000 bp) and the cDNA of *Plcd4* (an HA tag was fused to the N-terminus of PLCD4). The PLCD4 was also observed along the kinocilia, with a significant coimmunostaining with α-tubulin, thus suggesting the formation of a signaling complex with mGlu2 and Gq (Figures 7D, 7E and S9F).

**Fig. 7.**
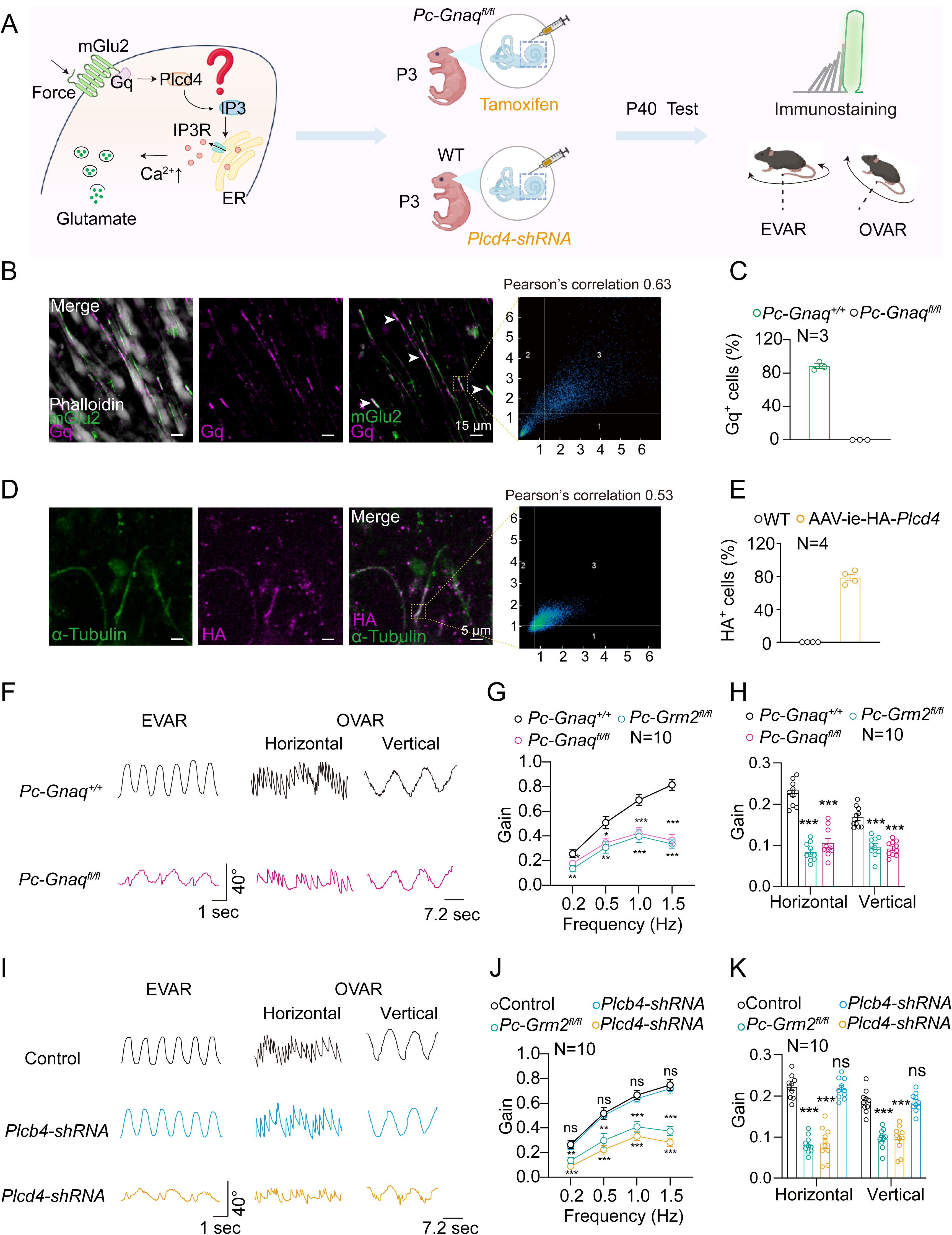
Gq and PLCD4 are required for normal balance. **(A)** Schematic diagram showing the research flow for exploring the vestibular functions of Gq and PLCD4. The *Pou4f3-CreER^+/−^*; *Gnaq^fl/fl^* mice (referred to as *Pc-Gnaq^fl/fl^*) mice were treated with tamoxifen to induce specific Gq ablation in hair cells, whereas the WT mice were injected with AAV-ie-*Plcd4-shRNA* through round window membrane to induce a specific knockdown of Plcd4 expression in the inner ear. The immunostaining and vestibular behavior tests were performed at P40. **(B)** Expression of mGlu2 (green) with Gq (magenta) and phalloidin (gray) of hair cells in utricle whole mounts of P40 *Pc-Gnaq^+/+^* mice. Arrows indicate coimmunostaining of mGlu2 with Gq. Scale bar, 15 μm. The Pearson’s correlation analysis of the fluorescence intensities of mGlu2 and Gq at the kinocilium was performed, revealing a correlation coefficient of 0.63. **(C)** Quantitative analysis of Gq expression in mGlu2-expressing utricular hair cells from *Pc-Gnaq^+/+^* mice or *Pc-Gnaq^fl/fl^* mice. Data are correlated to Fig. 7B (N = 3 mice per group). **(D)** Coimmunostaining of PLCD4-HA (magenta) with α-tubulin (green) of hair cells in utricle whole mounts of P40 AAV-ie-HA*-Plcd4* treated mice. Scale bar, 5 μm. The Pearson’s correlation analysis of the fluorescence intensities of α-tubulin and PLCD4 at the kinocilium of utricular hair cells was performed, revealing a correlation coefficient of 0.53. **(E)** Quantitative analysis of PLCD4 expression in utricular hair cells from WT mice or AAV-ie-HA*-Plcd4* treated mice. Data are correlated to Fig. 7D (N = 4 mice per group). **(F)** Representative recording curves of the VOR responses of *Pc-Gnaq^+/+^* mice and *Pc-Gnaq^fl/fl^* mice to earth-vertical axis rotation or off-vertical axis rotation. **(G, H)** Quantification of the VOR gain response of *Pc-Gnaq^+/+^* mice, *Pc-Gnaq^fl/fl^* mice and *Pc-Grm2^fl/fl^* mice to earth-vertical axis rotation **(G)** or off-vertical axis rotation **(H)** (N = 10 mice per group). **(I)** Representative recording curves of the VOR responses of WT mice, *Plcb4-shRNA* mice and *Plcd4-shRNA* mice to earth-vertical axis rotation or off-vertical axis rotation. **(J, K)** Quantification of the VOR gain response of WT mice, *Plcb4-shRNA* mice, *Plcd4-shRNA* mice and *Pc-Grm2^fl/fl^* mice to earth-vertical axis rotation **(J)** or off-vertical axis rotation **(K)** (N = 10 mice per group). (**G, H, J, K**) *P < 0.05; **P < 0.01; ***P < 0.001; ns, no significant difference. Gene knockout or knockdown mice compared with WT mice. The bars indicate mean ± SEM values. Data were statistically analyzed using one-way (**H, K**) or two-way (**G, J**) ANOVA with Dunnett’s post hoc test.

We next generated hair cell-specific Gq KO mice by crossing *Gnaq^fl/fl^* mice with *Pou4f3-CreER^+/−^* mice, and inner ear-specific *Plcd4*-knockdown mice via delivery of AAV-ie-*Plcd4*-*shRNA* through round window membrane injection (Figures 7A, S9A and S9B). Specific ablation of Gq or knockdown of PLCD4 in peripheral hair cells, but not in the central nervous system, such as in the vestibular nuclei in the brainstem, was verified by both western blotting and immunostaining (Figures 7B-7E, S8D and S9C-S9D). We measured the VORs of *Gnaq*-deficient or *Plcd4*-knockdown mice at P40. Notably, *Pou4f3-CreER^+/−^*; *Gnaq^fl/fl^* mice presented similar VORs as *Pou4f3-CreER^+/−^*; *Grm2^fl/fl^*mice in response to both earth-vertical and off-vertical rotation, with an approximately 30%-60% reduction in compensatory VOR gain in *Pou4f3-CreER^+/−^*; *Gnaq^fl/fl^* mice compared with their control littermates (Figures 7F-7H). Similarly, the mice treated with AAV-ie-*Plcd4-shRNA* presented decreased horizontal and vertical VOR gains in response to off-vertical rotation, which were comparable to those of *CreER^+/−^*; *Grm2^fl/fl^* mice (Figures 7I and 7K). Intriguingly, compared with *CreER^+/−^*; *Grm2^fl/fl^* mice, AAV-ie-*Plcd4-shRNA*-treated mice presented slightly but not significantly greater impairment of VORs in response to earth-vertical rotation, which might reflect a mGlu2-independent regulatory role for *Plcd4* in vestibular functions (Figures 7I and 7J). As a negative control, AAV-ie-*Plcb4-shRNA*-treated mice behaved normally in all VOR tests, which was consistent with the dispensable role of *Plcb4* in regulating mGlu2-mediated Ca^2+^ responses upon mechanical stimulation (Figures 7I-7K). Collectively, these results further support an essential balance-regulating role of the Gq-PLCD4 pathway in VHCs.

### mGlu2-Gq axis regulates TMC-independent mechanochemotransduction in VHCs

Our observation that mGlu2 expressed at kinocilia can regulate MET-independent Ca^2+^ signaling in VHCs prompted us to investigate the contribution of this Gq-mediated pathway to mechanical force-stimulated total Ca^2+^ responses. Recent evidence from multiple lines of investigation suggests that TMC1 and TMC2 are the probable pore-forming subunits of the MET channel complex at the tips of stereocilia^19–21^. Accordingly, simultaneous knockout of *Tmc1* and *Tmc2* in mice completely abolished MET currents, which could theoretically block mechanical force-stimulated Ca^2+^ influx via cation channels^22^. We tested this possibility by recording the Ca^2+^ signals of utricular hair cells in response to fluid jet stimulation and revealed that the fluid jet-induced Ca^2+^ response was reduced by approximately 80% in utricular hair cells from *Tmc1^−/−^; Tmc2^−/−^*mice compared with those from WT mice (Figures 8A-8C). Interestingly, the remaining 20% of the Ca^2+^ response was further decreased by 75% when treating the cells with mGlu2 inhibitor JNJ-40411813, which effectively blocked mechanical force-induced Gq activation downstream of mGlu2 (Figures 8A-8C and S9G). Moreover, pretreatment of *Tmc1^−/−^; Tmc2^−/−^* utricular hair cells with the Gq inhibitor YM-254890 completely abolished the residual Ca^2+^ response (Figures 8D and 8E). In agreement with these data, the application of YM-254890 alone to WT hair cells led to an approximately 20% reduction in the fluid jet-induced Ca^2+^ response (Figures 8D and 8E). Therefore, the mGlu2-Gq axis regulates TMC-independent mechanochemotransduction in VHCs, which accounts for approximately 15–20% of the mechanical force-induced Ca^2+^ response.

**Fig. 8.**
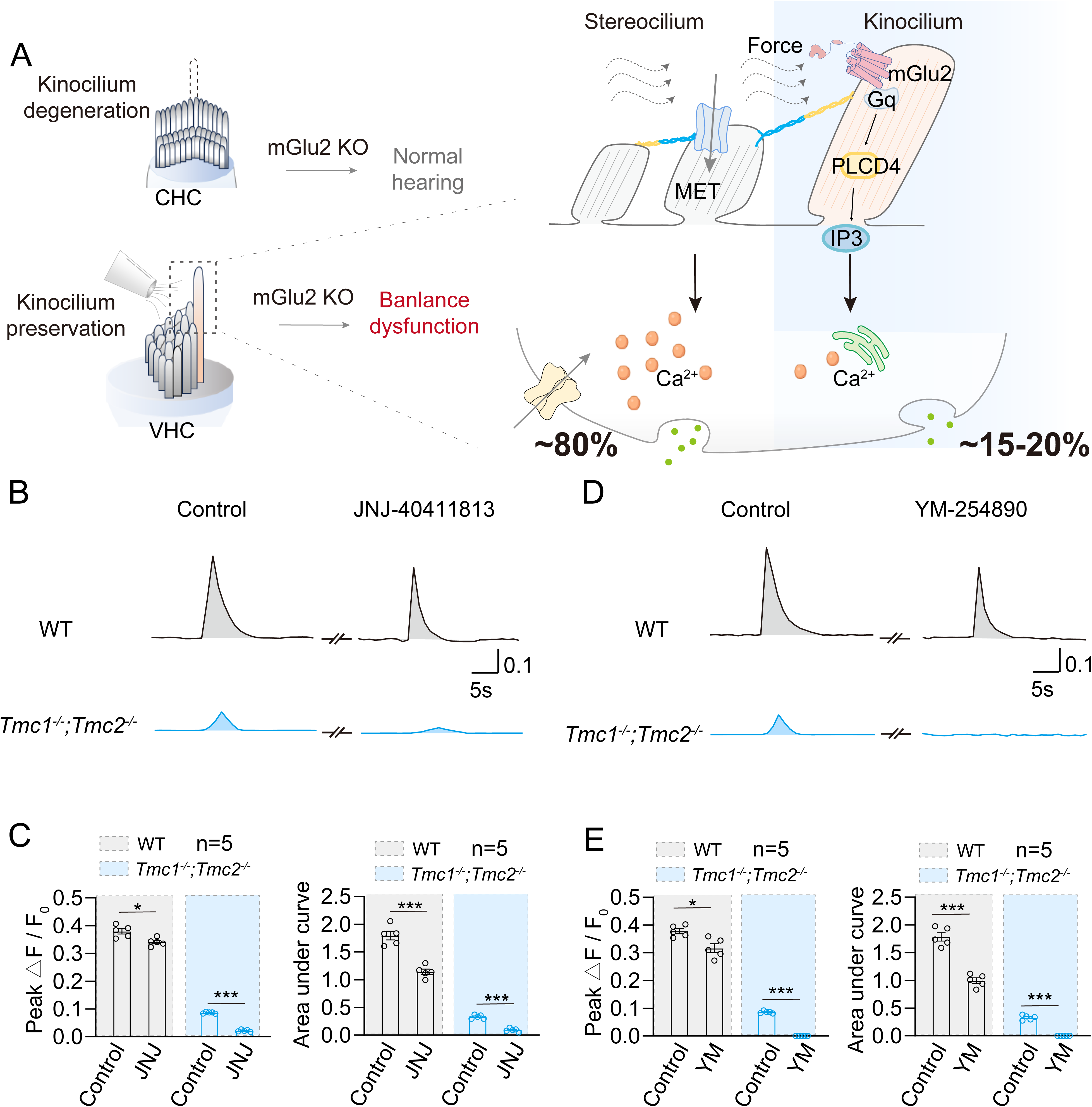
mGlu2 mediates TMC-independent mechanochemical transduction in VHCs. **(A)** Schematic illustration showing the fluid jet-stimulated Ca^2+^ responses in utricular hair cells via both MET process and a mGlu2-Gq-PLCD4 pathway in kinocilium. **(B, C)** Representative traces (**B**) and quantitative analysis (**C**) of fluid jet-stimulated Ca^2+^ signals in utricular hair cells in the absence or presence of 50 μM JNJ-40411813 derived from WT (black) or *Tmc1^−/−^; Tmc2^−/−^* mice (blue) (n=5). The magnitude of the Ca^2+^ response was characterized by ΔF/F_0_. **(D, E)** Representative traces (**D**) and quantitative analysis (**E**) of fluid jet-stimulated Ca^2+^ signals in utricular hair cells in the absence or presence of 5 μM YM-254890 derived from WT (black) or *Tmc1^−/−^; Tmc2^−/−^* mice (blue) (n=5). The magnitude of the Ca^2+^ response was characterized by ΔF/F_0_. **(C, E)** *P < 0.05; ***P < 0.001. Utricular hair cells pretreated with JNJ-40411813 or YM-254890 with those treated with control vehicle. The bars indicate mean ± SEM values. All data were statistically analyzed using unpaired two-sided Student’s t test.

## Discussion

### Mechanosensitive GPCR-second messenger system contribute to neurotransmitter release in VHCs

The traditional view regarding the role of VHCs in equilibrioception focuses on MET process at stereocilia, which converts mechanical stimuli into electrical signals via MET channels, followed by an increase in intracellular Ca^2+^ signaling and the release of neurotransmitters to convey motion information from the head to the central nervous system. Our current results indicate that a GPCR expressed in nearly 90% of VHCs, which belongs to the class C GPCR family, is able to sense force and induces an increase in intracellular Ca^2+^ concentrations and neurotransmitter release through the Gq-PLCD4 pathway. Notably, approximately 20% of the fluid jet-stimulated Ca^2+^ signal remaining in mGlu2-positive VHCs from TMC1/TMC2 double KO mice could be blocked by the mGlu2 antagonist JNJ-4041813 or the Gq inhibitor YM-254890. Moreover, VHC-specific deficiency of either mGlu2 or Gq, or knockdown of PLCD4 expression in VHCs impaired balance in mice and significantly decreased neurotransmitter release. These findings collectively suggest that the mechanosensitive GPCR-second messenger system in VHCs contributes to normal balance.

### Important functional role of kinocilia in VHCs

Unlike mature cochlear hair cells, mature VHCs retain their kinocilia, the only true cilia in hair cells^23^. Conventionally, since the MET in VHCs is thought to be regulated by the deflection of stereocilia, the kinocilia seem to be dispensable for equilibrioception. However, the functional role of kinocilia in normal balance remains largely unclear^24,25^. Here, we reveal that the mechanosensitive receptor mGlu2 is specifically expressed at the kinocilia of VHCs throughout life and is necessary for normal balance but not required for hearing. The activation of mGlu2 is involved in mechanical force-induced increases in intracellular Ca^2+^ concentrations and neurotransmitter release. Deficiency of mGlu2 in VHCs in *Pou4f3-CreER^+/−^*; *Grm2^fl/fl^* mice impaired the balance of mice without affecting the morphology of the utricular macula or the MET machinery in hair cells, whereas reintroduction of mGlu2 specifically in the VHCs of mGlu2-deficient mice restored normal balance. Moreover, our data suggest that downstream of mGlu2 activation, both Gq and PLCD4, key components of the Gq signaling pathway, are expressed in kinocilia. Therefore, our results indicate that a MET-independent system consisting of mechanosensitive GPCRs, Gq and PLCD4 is present in kinocilia and is required for the maintenance of normal balance, thus providing evidence of the functional importance of kinocilia in mature VHCs.

### Limitations of the study

The kinocilia in utricular hair cells are oriented in opposite directions on either side of the line of polarity reversal (LPR), which separates the utricle macula into two regions with opposite hair bundle orientations. Although our results indicate that mGlu2 is expressed in approximately 90% of utricular hair cells, the quality of the mGlu2 immunostaining results was not sufficient to provide precise information regarding the orientation of mGlu2 on individual kinocilia. Therefore, further in-depth investigations using super-resolution techniques is required to determine the precise expression pattern of mGlu2 on kinocilia, which may provide insights into whether mGlu2 modulates positional information and conveys specific balance-related information to the brain. In addition, while the Gq inhibitor YM-254890 almost completed eliminated the calcium signal in the utricular hair cells of *Tmc1^−/−^; Tmc2^−/−^* mice, a residual calcium signal was detected when we administered a mGlu2 antagonist (JNJ-40411813) in the fluid jet assay. Therefore, there might be other GqPCRs expressed in VHCs that potentially participate in mechanosensory transduction.

Our current study suggests that mGlu2 expression on kinocilia mediates mechanosensory transduction through the Gq pathway and generates calcium signals independent of MET at stereocilia. The MET-induced increase in the intracellular calcium concentration starts within a few milliseconds^26^, whereas mGlu2-mediated calcium signaling has a delay of less than 500 ms, suggesting that these processes may contribute to balance perception at different temporal scales. The physiological significance of these two pathways in balance perception remains to be further explored. To define PLCD4 localization, the AAV-delivered overexpression approach has been used. The use of PLCD4 knock-in mice with the desired tag could provide better resolution and more reliable information. Clarifying the expression patterns of mGlu2-Gq-PLCD4 axis components on kinocilia could help us better understand the potential compartmentalized GPCR signaling.

Taken together, our studies reveal that a mechanosensitive class C GPCR, mGlu2, is expressed at the kinocilia of VHCs and is required for normal balance. Mechanosensation by mGlu2 activates Gq-PLCD4 signaling, increases the intracellular calcium concentration and promotes neurotransmitter release. The discovery of the functional importance of the previously uncharacterized GPCR-Gq-PLCD4 signaling axis in kinocilia broadens our knowledge of balance sensation, one of the most important senses.

## Data availability

All the data generated in this study are included in the main text and/or supplemental information.

## Acknowledgments

We would like to thank Prof. Zhaofa Wu for sharing the glutamate sensor, and Prof. Yulong Li for providing suggestions on neurotransmitter detection. This work was supported by the National Science Fund for Distinguished Young Scholars Grant (82425105 to J.-P.S., 82225011 to X.Y.), the Foundation for Innovative Research Groups of the National Natural Science Foundation of China (T2321004 to J.-P.S.), the National Key R&D Program of China (2023YFA1801100 and 2024YFA1107500 to Z.Y.), the National Science Fund for Excellent Young Scholars (82422072 to Z.Y.), the National Natural Science Foundation of China (32130055, 32361163612 and 82330118 to J.-P.S., 92357303 to X.Y., 82271176 to W.-W.L., 82400838 to M.-W.W., 82501400 to J.-N.X.), the Taishan Scholars Program of Shandong Province (tsqn202408028 to Z.Y., tsqn201909189 to W.-W.L.), the Shandong Provincial Natural Science Fund (ZR2023QH394 to M.-W.W.). J.-P.S. is also supported by the Tencent New Cornerstone Science Foundation and Open Research Project in State Key Laboratory of Vascular Homeostasis and Remodeling (Peking University).

## Author contributions

J.-P.S., Z.Y., Y.S. and X.Y. initiated, designed and supervised the overall project. J.-P.S., Z.Y. and X.Y. started the screening of mechanosensitive GPCRs in vestibular systems. Y.S. and R.-J.C. provided guidance and suggestions on vestibular and cochlear studies. Z.Y. and J.-P.S. developed the magnetic beads assay. X.Y. designed and supervised the electrophysiology experiments and Ca^2+^ imaging. S.-H.Z., X.-H.W., Q.-Y.Z. and W.-F.Z. performed cell-based assays. S.-H.Z., Q.-Y.Z. and X.-H.W. performed electrophysiological experiments and Ca^2+^ imaging. S.-H.Z., J.-N.X and W.-F.Z. performed the glutamate detection experiments. J.-P.S., Y.S, Z.Y. and J.-L.W. participated in the design of gene knockout mice and modified AAV-ie. S.-H.Z., X.-H.W., Q.-Y.Z. and J.-N.X. performed vestibular behavior studies. S.-H.Z., X.-H.W., Q.-Y.Z., W.-F.Z., M.-W.W., and Y.-Q.W. performed immunofluorescence studies and ex vivo experiments. S.-H.Z., and Q.-Y.Z. performed single-cell qRT-PCR. S.-H.Z., X.-H.W., Q.-Y.Z. and J.-H. D performed western blotting analyses. X.-H.W. and Q.-Y.Z. performed AAV-ie-related experiments. J.-P.S., Y.S., X.Y., Z.Y., S.-H.Z., X.-H.W., Q.-Y.Z., J.-N.X. and J.-L.W. participated in data analysis and interpretation. Y.S., P.H. and W.-W.L. provided insightful ideas related to equilibrioception based on clinical practice. S.-H.Z., X.-H.W., Q.-Y.Z. and J.-N.X. prepared the figures. J.-P.S., Z.Y., Y.S. and X.Y. wrote the manuscript. All the authors have read and commented on the manuscript.

## Competing interests

The authors declare no competing interests.

## Notes

### Competing Interest Statement

The authors have declared no competing interest.

